# Bringing TB genomics to the clinic: A comprehensive pipeline to predict antimicrobial susceptibility from genomic data, validated and accredited to ISO standards

**DOI:** 10.1101/2023.11.04.565651

**Authors:** Kristy A Horan, Linda Viberg, Susan A Ballard, Maria Globan, Wytamma Wirth, Katherine Bond, Jessica R Webb, Thinley Dorji, Deborah A Williamson, Michelle L Sait, Ee Laine Tay, Justin T Denholm, Benjamin P Howden, Torsten Seemann, Norelle L Sherry

**Affiliations:** Microbiological Diagnostic Unit Public Health Laboratory (MDU-PHL), Department of Microbiology & Immunology, University of Melbourne at the Peter Doherty Institute for Infection & Immunity, Melbourne, Victoria, Australia; Mycobacterium Reference Laboratory, Victorian Infectious Diseases Reference Laboratory (VIDRL), at the Peter Doherty Institute for Infection and Immunity, Melbourne, Victoria, Australia; Department of Microbiology, Royal Melbourne Hospital, Melbourne, Victoria, Australia; Department of Microbiology and Immunology, The University of Melbourne, at the Peter Doherty Institute for Infection and Immunity, Melbourne, Victoria, Australia; Centre for Pathogen Genomics, The University of Melbourne, Melbourne, Victoria, Australia; School of Biological Sciences, University of Adelaide, Adelaide, Australia; Centre for Epidemiology and Biostatistics, Melbourne School of Population and Global Health, The University of Melbourne, Melbourne, Victoria, Australia; Communicable Disease Epidemiology and Surveillance, Health Protection Branch, Public Health Division, Department of Health, Victoria, Australia; Victorian Tuberculosis Program, Melbourne Health, at the Peter Doherty Institute for Infection and Immunity, Melbourne, Victoria, Australia; Department of Infectious Diseases, The University of Melbourne, at the Peter Doherty Institute for Infection and Immunity, Melbourne, Victoria, Australia; Department of Infectious Diseases & Immunology, Austin Health, Heidelberg, Victoria, Australia

## Abstract

**Background:** Whole genome sequencing (WGS) is increasingly contributing to the clinical management of tuberculosis. Whilst the availability of bioinformatic tools for analysis and clinical reporting of *Mycobacterium tuberculosis* sequence data is improving, However, there remains a need for accessible, flexible bioinformatic tools that can be easily tailored for clinical reporting needs in different settings and are suitable for accreditation to international standards.

**Methods:** We developed tbtAMR, a flexible yet comprehensive tool for analysis of *Mycobacterium tuberculosis* genomic data, including inference of phenotypic susceptibility and lineage calling. Validation was undertaken using local and publicly-available real-world data (phenotype and genotype) and synthetic genomic data to determine the appropriate quality control metrics and extensively validate the pipeline for clinical use.

**Findings:** tbtAMR accurately predicted lineages and phenotypic susceptibility for first- and second-line drugs, with equivalent computational and predictive performance compared to other bioinformatics tools currently available. tbtAMR is flexible with modifiable criteria to tailor results to users’ needs.

**Interpretation:** The tbtAMR tool is suitable for use in clinical and public health microbiology laboratory settings, and can be tailored to specific local needs by non-programmers. We have accredited this tool to ISO standards in our laboratory, and it has been implemented for routine reporting of AMR from genomic sequence data in a clinically relevant timeframe. Reporting templates, validation methods and datasets are provided to offer a pathway for laboratories to adopt and seek their own accreditation for this critical test, to improve the management of tuberculosis globally.

**Funding:** Victorian Government Department of Health; Australian National Health and Medical Research Council and Medical Research Futures Fund.

**Research in context:** *Evidence before this study:* We searched PubMed for studies using the search terms: “Mycobacterium tuberculosis”, “clinical”, “bioinformatics”, “genomics”, “drug resistance (OR antimicrobial)”, published prior to 31^st^ July 2024 without language restrictions (n=258). We considered all studies from this search that used genomics to infer likely drug-resistance in *M. tuberculosis*. Many of these studies highlight the challenges in making meaningful interpretations from genomic data for the purpose of inferring AMR for clinical applications. Despite the development of bioinformatics tools and compilation of catalogues of resistance conferring mutations, few of these studies directly address the challenges specific to implementation of a whole genome sequencing (WGS) and bioinformatics pipelines to deliver validated and accredited results for use in clinical applications and none provide sustainable solutions.

*Added value of this study:* To the best of our knowledge, this study is the first to detail practical solutions to challenges to validating and accrediting the routine inference of AMR from WGS data for clinical applications for treatment of *M. tuberculosis*. We developed, validated and accredited a bioinformatics tool, tbtAMR, which is robust and flexible, using a data-driven approach to inferring resistance, allowing for simple customisation of behaviour on a large collection of in-house data supplemented with publicly available data. Furthermore, we have made this pipeline and the accompanying data available for use by the wider public health community to aid others in implementing similar programs. Consultation with infectious disease clinicians, microbiologists and epidemiologists has allowed us to develop a robust workflow, that encompasses sequencing, analysis, interpretation, reporting, discrepancy resolution and ongoing verification.

*Implications of all the available evidence:* This study addressed the specific needs of reporting genomic AMR for *M. tuberculosis* in a clinical and public health laboratory. Furthermore, this work makes available a pipeline and data for laboratories to utilise to implement genomic AMR for *M. tuberculosis*. Through the use and further development of publicly available and open-source bioinformatics software, we have demonstrated the feasibility of clinical reporting of genomic AMR results for *M. tuberculosis* in a low-incidence high-income setting. We believe that this work lays the foundation for making reporting of genomic AMR for *M. tuberculosis* more accessible to public health and reference laboratories in general.

## Introduction

*Mycobacterium tuberculosis* (Mtb) is the causative agent of tuberculosis (TB), a disease that is globally distributed but predominantly affecting people in low- and middle-income communities and is the leading cause of death from a single infectious agent.^1^

Treatment of TB disease is lengthy and expensive, requiring combination therapy with at least two antimycobacterial drugs prescribed for a minimum of six months.^2,3^ This is usually commenced with four antimycobacterial drugs (‘first-line agents’): rifampicin, isoniazid, pyrazinamide and ethambutol, followed by rationalisation to two agents (rifampicin and isoniazid, if susceptible). If drug resistance is detected, an alternative regimen including second- and third-line agents will be used (Supplementary Figure 1 and Table 1).

The slow growth of Mtb *in vitro* means that clinicians treating patients with TB must choose treatment regimens before phenotypic drug susceptibility test (DST) results are available, which can take up to two months^4^ in many settings. Whole genome sequencing (WGS) has the advantage of providing comprehensive results from an Mtb isolate in a single test (rather than individual tests for species identification, lineage, and drug susceptibility), potentially significantly reducing turnaround time to DST for both first-line and other antimycobacterial drugs, especially when undertaken from Mycobacteria Growth Indicator Tube (MGIT) culture. In addition, WGS can provide increased reliability of resistance detection for some drugs, due to known challenges with phenotypic reproducibility, such as ethambutol and pyrazinamide^5^. WGS data is also being used to provide support for outbreak investigation and contact tracing^6^ (Figure 1A). Therefore, the use of routine WGS for Mtb has the potential to improve patient management in a clinical setting and public health outcomes, and new Australian national guidelines recommend routine use and reporting of findings for these reasons^7^.

**Figure 1.**
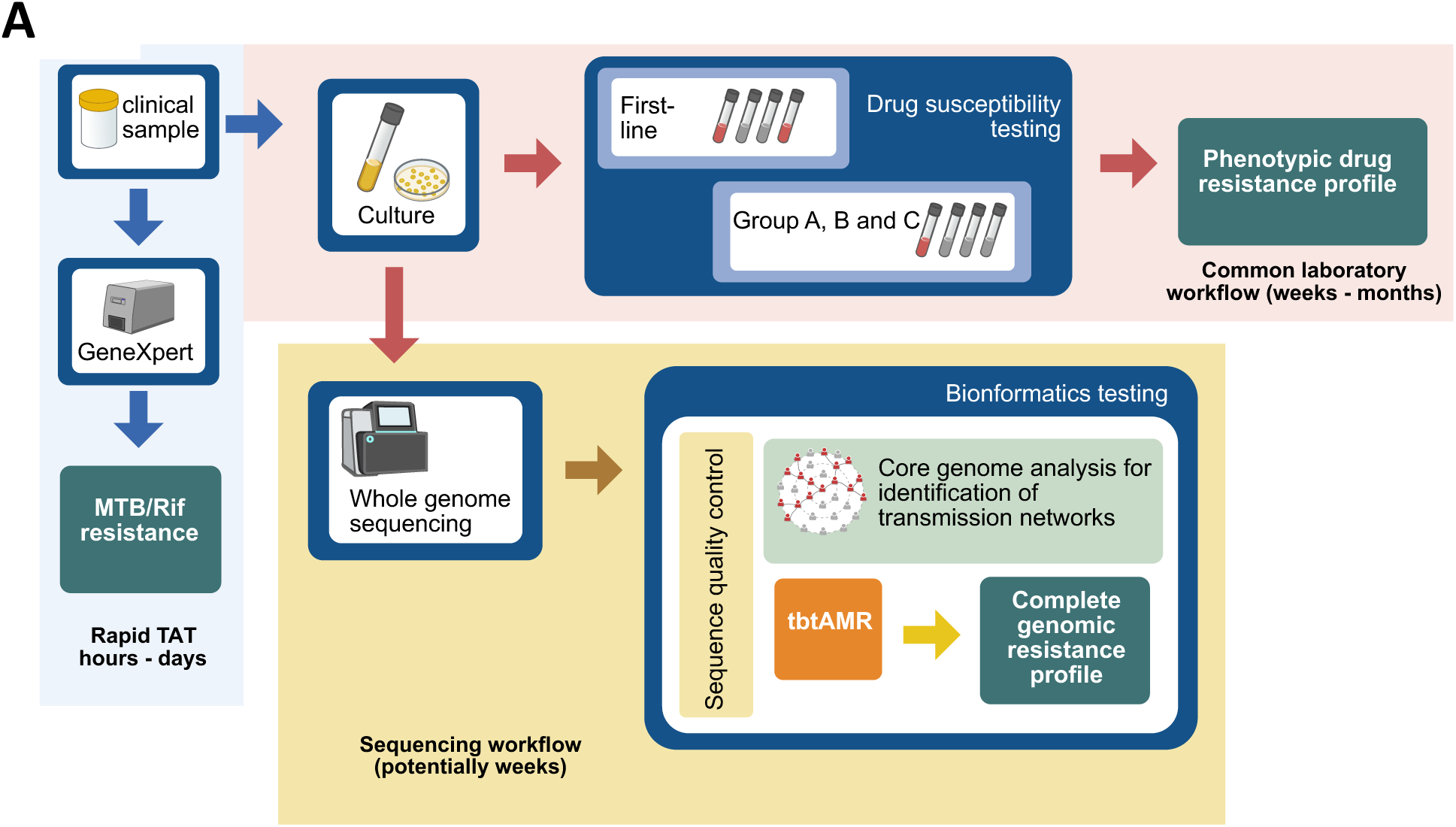
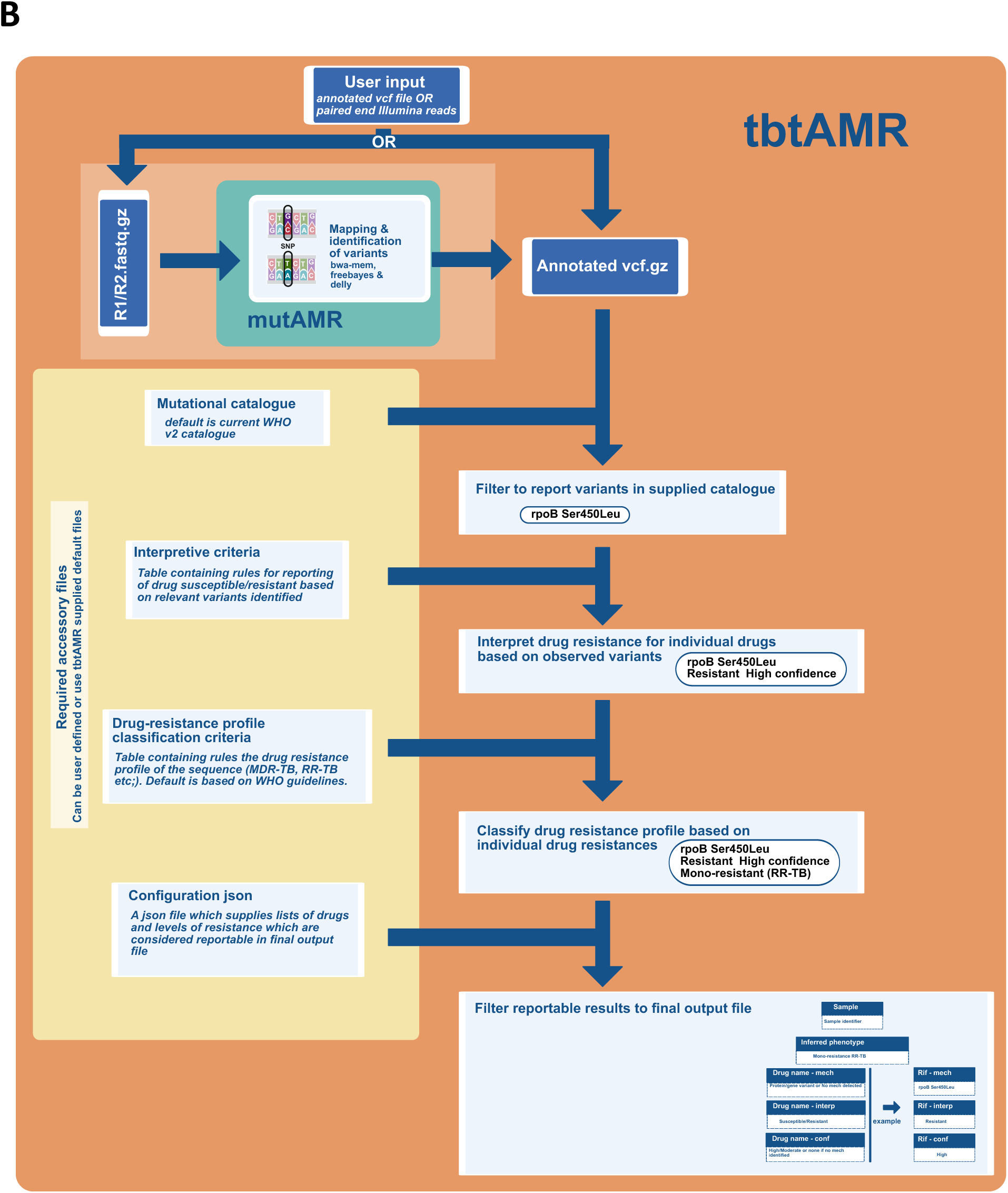
Incorporation of tbtAMR into *M. tuberculosis* diagnosis and testing workflow in a reference laboratory. **A. Diagnostic testing pathways for Mtb:** *M. tuberculosis* and potential rifampicin resistance can be identified rapidly using GeneXpert (closed-system PCR for most common rifampicin resistance mutations) in a diagnostic laboratory (turnaround time hours–days). Traditional culture methods require samples to be inoculated to solid media (agar slopes) and/or liquid media (most commonly MGIT). Cultures may take up to 6-8 weeks to flag positive. From positive culture, phenotypic drug susceptibility testing is then commenced, taking up to two weeks to be completed, or longer if cascade testing is used (Group A, B and C drugs only set up when resistance to first-line drugs is identified). Total turnaround times from sample to culture and phenotypic DST results is usually 6 weeks-3 months. If optimised, whole genome sequencing can potentially reduce this turnaround time and provide a comprehensive resistance profile without the need for further testing, whilst also providing important contributions to public health control efforts through contact tracing. **B. tbtAMR pipeline:** tbtAMR inputs can be either an annotated variant file (vcf format, from short-read or long-read sequences) based on H37rV v3 reference genome or paired-end short-read data (fastq file), making tbtAMR sequencing platform agnostic. tbtAMR requires 4 accessory files, defaults are supplied with tbtAMR, but users can their own if appropriate. By default tbtAMR uses version 2 of the WHO mutational catalogue as default with accompanying files containing criteria for interpretation, classification and reporting. The accessory files are the mechanism by which tbtAMR provides interpretation of detected variants, performs classification of drug resistance profile and filters results for reporting in final output file. Abbreviations: Mtb, *Mycobacterium tuberculosis*; Rif, rifampicin; TAT, turnaround time. Variant nomenclature example: “rpoB_Ser450Leu” denotes a change in the amino acid from serine to leucine at position 450 in the RpoB protein sequence.

Despite the urgent need for routine identification of Mtb AMR from genomic data to inform therapy and the wealth of available bioinformatic resources, few clinical and public health laboratories (CPHLs) globally have processes in place to routinely report AMR from WGS data for Mtb. Many factors need to be considered when implementing such an AMR detection program, particularly where patient management may be impacted, including performance, ease-of-use and flexibility of tools in predicting drug-resistance, the impact of laboratory methodologies and existing processes, such as existing bioinformatics pipelines, reporting structure, CPHL data management, and test accreditation by national accreditation bodies to International Organization for Standardization (ISO) standards (ISO15189:2022).^8^ Laboratories seek accreditation to ISO standards to demonstrate that they are able to produce robust, reproducible and high-quality laboratory services; the process of obtaining accreditation, and the difficulties in accrediting genomic tools, is covered in depth elsewhere^9^.

There are many high-quality and established bioinformatics tools available for identification of genomic determinants of AMR in Mtb, including Mykrobe,^10^ TB-Profiler^11^ and ariba^12^, with others being recently evaluated^13^. Additionally, a second version of the WHO Mtb Mutation Catalogue^10,14^ has now been released with an expanded catalogue of mutations for second- and third-line drugs (Groups A-C Table 1), as well as guidance for interpreting complex genomic interactions. Whilst this additional information is welcome and informative for detailed analysis, clinical reporting often requires a more pragmatic approach as clinicians may find this commentary difficult to interpret, preferring more binary results (Resistant/Susceptible) where possible. For reasons such as this, users may wish to modify the protocols, or ‘rules’, for interpretation of each variant. Alternatively, they may wish to add other mutations of local interest into the resistance database used for analysis, or change the way that mutations are classified (for example, swapping ‘Susceptible’ for ‘Wildtype’) according to their local needs. Most of the currently used tools for Mtb genomic DST (gDST) do not currently allow users to make similar changes without a bioinformatician to modify existing code, limiting the ability of clinical and public health labs, as well as researchers, to adapt tools to meet local needs. Furthermore, in high-throughput sequencing labs, WGS may be undertaken in large batches with existing workflows and pipelines in place, therefore bioinformatics tools need to be able to be easily incorporated into these existing pipelines, be straightforward to maintain and for any updates to be assessed easily by reverification.

Here we describe the validation of a real-time routine analysis tool for inference of phenotypic resistance in Mtb, based on detection of genomic variation, in a clinical and public health microbiology setting. We have subsequently gained accreditation to ISO standard 15189:2022 (Clinical Laboratories) for this tool, tbtAMR, for clinical and public-health reporting of genomic AMR detection in Mtb. The pipeline and validation approach are made available here for other laboratories globally to aid in the transition to WGS for genomic AMR reporting.

## Methods

### tbtAMR design

Many software options exist for detection of AMR determinants in Mtb sequences, including Mykrobe^10^ and TBProfiler^11^, each with pros and cons. When developing a stand-alone tool, we considered that the process must be easy to incorporate into existing laboratory workflows (Figure 1A), have flexibility in key outputs, and to be easily maintained to expedite any updates, ideally by non-developers. These were noted limitations of existing tools for use in a CPHL, where software maintenance and updates can be challenging. In addition, reporting of AMR from WGS data for Mtb will directly impact patient management. Thus, the results generated should be suitable for reporting in a clinical setting with a layer of interpretation that is fit-for-purpose, not reporting raw (unfiltered) results unless required for a specific purpose. In tbtAMR, we report four key parameters to assist in clinical decision making: (i) presence or absence of mutations including small insertions or deletions (indels), large structural changes, gene loss (or loss of function) and SNPs, (ii) confidence that this mutation is associated with drug resistance, (iii) the level of drug resistance conferred by this mutation (high or low), and (iv) the overall resistance profile for the isolate based on the resistance inferred to individual drugs (e.g. based on WHO definitions).

tbtAMR (https://github.com/MDU-PHL/tbtamr), is a pure python package and can be used as a stand-alone tool with minimal dependencies, taking an annotated variant file (vcf) to generate an inferred antibiogram and resistance profile based on user-defined criteria (Figure 1B). The recently updated WHO mutation catalogue for Mtb (version 2) is currently implemented as the default mutational catalogue within tbtAMR^14^ with its accompanying guidelines documentation^15^ informing criteria for classification. However, a user-defined catalogue of mutations can also be supplied, along with bespoke criteria for interpretation and classification according to a user’s needs, allowing for increased flexibility and ease of update.

Providing reportable interpretations of the likely resistance to each drug is a key requirement of tbtAMR. Unlike many approaches, tbtAMR does not rely on hard coded or installed library files to undertake this interpretation, but allows the user to choose between a default library and user-modified files. tbtAMR relies on four accessory files (detailed below and Figure 1B, and available at^16^): (i) a mutational catalogue or database of resistance mechanisms; (ii) rules for interpreting resistance (such as categorisation as susceptible/resistant for each drug); (iii) rules for reporting classification of resistance profile (such as ‘fully susceptible’ or ‘MDR-TB’); and (iv) a config file defining the structure of the mutational catalogue and report outputs.

- *Mutational catalogue* – a CSV file containing the drugs, genes, variants and confidence levels for calling drug resistance. The supplied default file is based on the WHO catalogue version 2^15^.

`o` *interpretation_criteria.csv* – this CSV file includes rules to interpret information from the mutational catalogue when a variant is detected. The supplied default interpretation rules file is based on the additional commentary supplied with the WHO mutational version 2 catalogue (see Supplementary File 1). There are two types of rules: ‘default rules’ applied broadly to most mutations, and ‘override rules’ for exceptions and edge cases. ‘Default rules’ describe rules for interpretation of reportable drug resistance in most circumstances; for example, for a certain drug, if a variant associated with resistance (or associated with resistance – interim) is present, then report ‘Resistant’ for the corresponding drug. Only drugs which have a default rule will be reported by tbtAMR, allowing the user to control which drugs are reported. ‘Override rules’ provide additional rules to describe interpretation of edge cases or exceptions to the default rules, hence overriding the default rule. For example, when a specific variant is detected, report “Low-level resistant” based on commentary in the WHO Catalogue. Users may supply additional rules, such as reporting a specific comment under certain conditions, or where the user wishes to report the presence of a variant but not an interpretation (examples in Supplementary Table 1).
- *classification_rules.csv* – these rules determine the overall classification profile, based on the combination of resistances in the sequence, for example mono-resistant or multidrug-resistant TB. The defaults are based on the current WHO guidelines (Supplementary Figure 2 and Supplementary File 1).
- *db_config.json* – this file provides information about the structure of the mutational catalogue, drugs to include, the values for confidence and resistance levels, and cascade reporting structure (more details in Supplementary File 1)(Figure 1B).

tbtAMR provides a default files for each of these accessory files, based on WHO catalogue of mutations version 2^15^ and the current WHO guidelines for classification of resistance profile^2^. Alternatively, users may choose to modify or replace these files with their own versions to suit their local requirements. As these are simple CSV files, these modifications may be made successfully by non-bioinformaticians, providing flexibility in interpretation and reporting. Note that tbtAMR performance has been investigated based on the default database and interpretation criteria; performance may vary where a bespoke database and/or interpretive criteria are used. The tool has optional additional functionalities, including paired-end read alignment, variant calling and annotation, which require additional dependencies including bwa-mem^17^, freebayes^18^, snpEff^19^, delly^20^, bcftools and samtools^21^ (Figure 1B). These additional functionalities are implemented in our locally-designed mutAMR tool (https://github.com/MDU-PHL/mutamr). The entire package is available as a conda package.

### Validation dataset

We included three data sources in our validation dataset (Supplementary Figure 2 and Supplemental File 2). Firstly, we included 2005 sequences generated at Microbiological Diagnostic Unit Public Health Laboratory (MDU PHL) from 2018–2022 and with phenotypic (DST) data generated at the Mycobacterial Reference Laboratory (MRL) (Supplementary Methods). Secondly, 13,777 publicly available sequences were included, selected to be representative of all major global lineages and with phenotypic data available for first-line agents and a subset also having data available Group A, B and Group C drugs^22–25^. Lastly, in order to assess the accurate detection of variants by tbtAMR, we also used simulated genomic data. Paired-end sequence data was generated using TreeToReads software^26–29^, directly from the H37rV reference strain (RefSeq accession NC_000962.3) or following introduction of 2–4 variants per 10,000 bases at known positions with an error-profile representative of the NextSeq500 instruments at MDU PHL (n= 1830). As resistance testing aims to identify resistant subpopulations of Mtb that may emerge during treatment, these simulated reads were mixed (Supplementary Figure 3) to simulate different allelic frequencies (0.01 to 0.99) across a range of average genome depth (20x to 200x – incremented by 20x).

### Measuring the accuracy of Mtb genomic sequence recovery

For simulated genomes, we tested the ability of the tool to correctly identify introduced variants using a minimum depth of 10X and 20X, meaning that to make a base call there was least 10 or 20 reads covering each position. Reads were aligned back to the reference genome and the correctness of base calling by mutAMR was assessed (Supplementary Figure 3).

### Analysis of phylogenetic lineage and inferred phenotypic DST from genomic data

We looked at how well inferred phenotype from detected genomic resistance mechanisms (single nucleotide polymorphisms [SNPs], insertions or deletions) correlated with phenotypic DST results. To quantify this, we compared the phenotypes inferred from the presence of SNPs, insertions or deletions in the genome sequences with actual phenotypic susceptibility profiles (Supplementary File 2), and calculated sensitivity, specificity, negative predictive value (NPV), positive predictive value (PPV) and accuracy recorded. When discrepancies between inferred-resistance and the reported phenotypic resistance are identified, these would ideally be resolved with additional phenotypic testing. However, since this was not possible for the discrepancies observed in the publicly available datasets, discrepancy resolution could not be undertaken and any such results were included as discrepancies in performance assessment. tbtAMR also determines phylogenetic lineage, including lineages 1–9 as well as animal adapted species/sub-species^11^. We also assessed the concordance between phylogenetic lineage determined by tbtAMR and the phylogenetic lineage that was supplied as part of the public dataset^25^ (Supplementary File 2).

### Ethics approval

Ethical approval was received from the University of Melbourne Human Research Ethics Committee (study number 1954615).

### Role of the funding sources

The funders played no part in data collection, analysis, interpretation, writing of the manuscript or decision to submit.

### Data access

KAH, TS, NLS, JW and TD had access to data; KAH, BPH, TS and NLS were responsible for the decision to submit the manuscript.

## Results

### Validation of variant detection

#### Accuracy of simulated sequence recovery by mutAMR

tbtAMR has the option to align and detect variants in paired-end Illumina WGS data, through mutAMR. We assessed the ability of mutAMR to recover known (introduced) SNPs in simulated sequences, and the minimum average sequence depth for reliable results. Most tools used for detecting variants use a minimum depth criteria of 10X^30,31^, meaning that there must be at least 10 reads aligned to a particular position in the reference genome. However, when we investigated SNPs identified across a range of average sequence depths and allelic frequencies, several false positive (FP) SNPs (mean 6.6, 95% confidence interval (CI) 3.9-9.3) were observed when using the standard criteria of 10X, which may result in false reporting of resistance mechanisms. Increasing the minimum depth for base calling to 20X improved performance, reducing the number of FP SNPs to a mean of 1.1 (CI 0.7-1.5), with little impact on the percentage of introduced SNPs recovered (Figure 2A-B). We further examined the impact of varying average genome depth and allelic frequency on the ability of mutAMR to accurately identify mutations in the simulated sequences. Where average genome depth was less than 40X and allelic frequency was less than 10%, sensitivity of variant identification was low at only 2.1% (CI 1.48%-2.66%) and False Discovery Rate (FDR) was 41.1% (CI 31.1%-51.2%). However, maintaining 20 reads minimum depth for base-calling, an average genome depth of ≥40X and allele frequencies ≥10% gave the best performance (Figure 2C-D), with an overall variant identification sensitivity of 96.3% (CI 95.5%-97.1%) and low FDR of 0.21% (CI 0.11%-0.31%). Detection of other variant types, such as deletions, by mutAMR was consistent with TB-profiler and Mykrobe, using publicly available data. Hence, the minimum acceptable read depth for identification of resistance conferring variants was set to ≥40X and minimum allele frequency of 10%.

**Figure 2.**
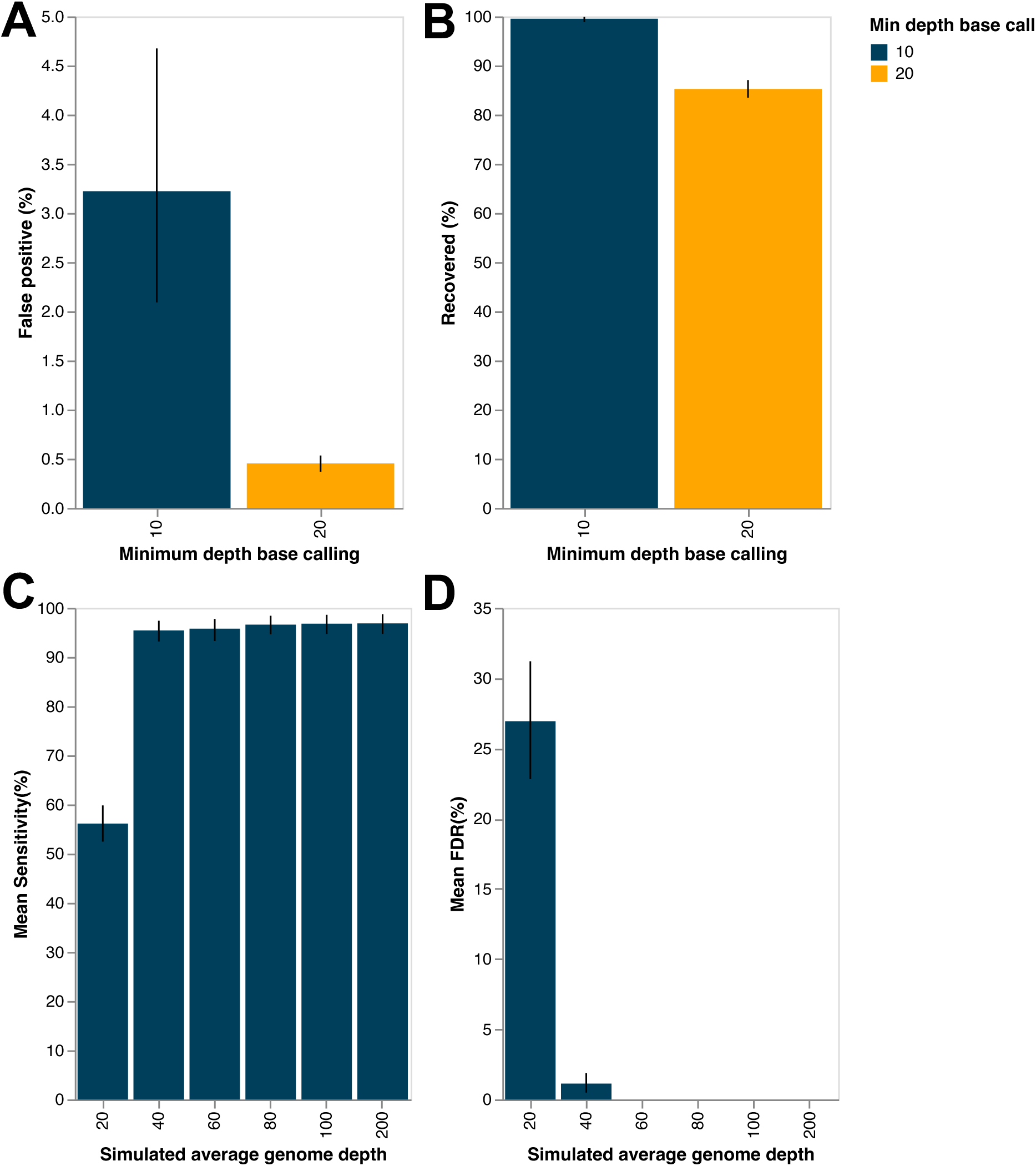
Performance of mutAMR to recover SNPs from Mtb sequences. To assess the performance of mutAMR, implemented by tbtAMR to accurately recover mutations in Mtb sequences, simulated genome data across a range of average genome depths from 10X to 200X at allelic frequencies 10-100% were assessed. The impact of minimum read depth for base calling at an individual position in the genome (depth 10 or 20) was assessed for: the percentage of false positive SNP calls (FP) identified (**Panel A**) and the percentage of introduced SNPs recovered by mutAMR (**Panel B**). Using a minimum-depth criteria of 20X for individual base calling, **C.** SNP calling sensitivity and **D.** false discovery rate (FDR) were assessed average genome depth using average genome depth of coverage of ≥40X and allelic frequency ≥10% (error bars indicate 95% confidence interval).

### Accuracy of phylogenetic lineage calling by tbtAMR

tbtAMR accurately identified lineages compared to lineages reported from public datasets^25^, (98.5% concordance 871/884). Of the 13 discordant results (Supplementary Table 2), 12 were cases where tbtAMR identified two different phylogenetic lineages in the sequence. In each case, the reported lineage was one of the lineages detected by tbtAMR. These could reflect mixed sequences or sequences where low coverage may impact base-calling at key sites that differentiate lineages from the reference lineage (lineage 4). It should be noted that the detection of a mixed lineage does not indicate that a sequence is of poor quality for detection of AMR variants.

### Inference of phenotypic AMR from genomic data

Many studies have explored the inference of phenotypic AMR from genomic data using various tools^10,24,32,33^. We do not aim to duplicate these high-quality studies in this study, but rather to assess the suitability of tbtAMR for inference and reporting of Mtb AMR to acceptable standards for clinical and public health purposes.

We began with a dataset of 15,323 sequences from both publicly available datasets^22–25^ and sequences generated in-house using sequencing workflows accredited to ISO standards. Implementing the quality control thresholds and limits of detection outlined above, we excluded 608 sequences based on quality metrics (Supplementary Figure 4), leaving 14,715 sequences of sufficient quality to include in the validation of tbtAMR for inference of phenotypic AMR.

The performance of tbtAMR in inferring phenotypic resistance to first-line drugs was excellent, with high performance across all first-line drugs (Table 2 and Figure 3) and was consistent with the performance of both TB-profiler and Mykrobe (Supplementary Figure 5). Lower positive predictive values for ethambutol (81.1% CI 79.4%-82.7%) were due to detection of variants in *embB* at codon 360 where phenotype reported was susceptible. This observation is likely due to known challenges in consistency of ethambutol DST and are consistent with other studies^10,24,32,33^. Prediction of pyrazinamide resistance also has lower sensitivity than other first-line drugs (80.0% CI 75.4%-84.0%) and is likely due both challenges in pyrazinamide DST and less robust understanding of the molecular mechanisms leading to pyrazinamide resistance^5^.

**Figure 3.**
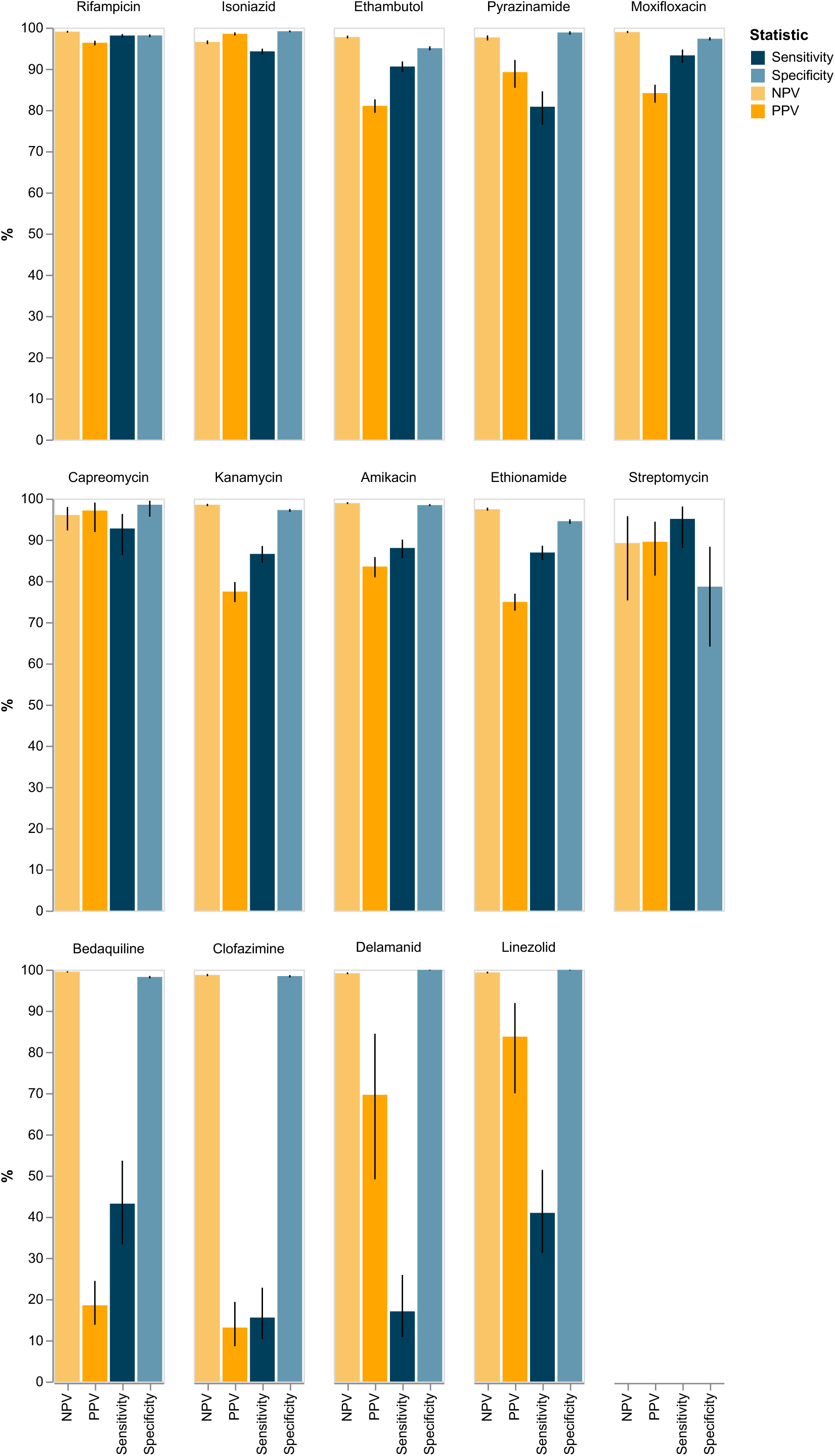
Performance of tbtAMR inference of phenotypic susceptibility. Inferred phenotypes predicted by the tbtAMR pipeline were compared to phenotypic DST data, true positive (TP) was recorded when a mutation was reported and the phenotype was resistant; a true negative (TN) was recorded when no mutation was reported and the phenotype was susceptible; a false positive (FP) was recorded when a mutation was reported, but the phenotype was susceptible, and a false negative (FN) was recorded when no mutation was reported, and the phenotype was resistant. Sensitivity, specificity, positive and negative predictive values calculated and expressed as a percentage (error bars indicate 95% confidence intervals). DST, direct susceptibility testing; PPV, positive predictive value; NPV, negative predictive value.

tbtAMR predicted resistance to groups A, B and C (Table 1) with performance comparable to other studies^10,24,32,33^ (Supplementary Figure). The high number of false-negative and false-positive results for prediction of phenotypic resistance (Table 2) are likely to be due to incomplete understanding of the mechanisms of resistance to these drugs, global paucity of data (phenotypic and genotypic) and challenging or inconsistent phenotypic testing for some drugs. Although tbtAMR performance was consistent with other tools and use of updated catalogue improved performance compared on the previous WHO catalogue, prediction of bedaqualine, delaminid, linezolid and clofazimine was not of sufficient performance to allow for routine reporting of resistance to these drugs from genomic data in a CPHL setting, due to the high rates of both false positive and false negative results (Supplementary Figure 6).

### Implementation process

As AMR genotypes and inferred antibiograms for Mtb were new to most clinicians, additional interpretive comments were created for inclusion in reports – for example, explaining the difference between confidence level and level of resistance in the inferred phenotype (Supplementary Methods and Supplementary Files 3). Clinical and laboratory colleagues were consulted about the draft report design to ensure the formatting and explanations were clear and unambiguous in our local setting. Once validation and report design were complete, Mtb genomic AMR inference using the tbtAMR tool was implemented into existing genomic workflows, including standard operating procedure documentation (SOP), staff training, incorporation into the laboratory information management system, and testing of reporting and feedback loops (e.g. reporting of samples that failed QC).

To address any discrepancies in phenotypic and genotypic results, a multidisciplinary expert panel for Mtb AMR was formed, meeting monthly to discuss individual cases and provide expert recommendations to clinicians where discrepancies were identified. This provides the opportunity for ongoing dialogue between the wet-lab microbiologists, clinicians, epidemiologists, medical microbiologists and bioinformaticians to monitor for AMR determinants of interest, discuss individual cases and discrepancies, and undertake prospective validation of novel or uncharacterised AMR determinants.

In preparation for ISO-equivalent accreditation, revalidation and reverification processes were also defined for tbtAMR (Supplementary Methods). Subsequently, the Mtb genomic AMR inference workflow has now been accredited to ISO 15189:2022 standards.

## Discussion

We have reported here the development of a custom Mtb AMR analysis and reporting pipeline for use in the clinical and public health microbiology setting, with easily customisable interpretative and classification criteria to maximise PPV and NPV for prediction of AMR in Mtb. Performance of inferred AMR using our process is consistent with the performance of other tools^10,33^ (Supplementary Figure 5). This pipeline has been robustly validated, implemented and accredited to ISO standards in our laboratory, with reverification strategies for ongoing improvement, and reporting processes put in place.

Globally, we have seen increasing availability of WGS in clinical and public health labs, particularly since the COVID pandemic, resulting in a massive increase in sequencing capacity and data^34^. However, significant skills gaps exist in the CPHL workforce^33,35,36^, particularly in LMICs, limiting the ability to use, develop and/or maintain open-source software, and proprietary alternatives may be expensive. Whilst open-source software is accessible and free to users, it can be challenging to implement in a CPHM laboratory setting. These tools are mostly developed and maintained by independent researchers, and may not always be maintained or frequently updated, may be difficult to install, or not meet the needs of the CPHM lab^37^. Maintaining and implementing such tools in an ISO-accredited laboratory structure can be time-consuming and difficult, requiring significant staff time and resources^9^, as laboratories cannot be selective about the updates adopted if the tool is maintained elsewhere, issues which are further exacerbated in resource-limited settings^38^. tbtAMR and its supporting documentation aims to address some of these issues for CPHM, by providing validation methods and datasets, reverification processes and example reports. Additionally, due to minimal additional dependencies, tbtAMR can easily be installed and run on a laptop or be incorporated into large and complex bioinformatics workflows easily.

tbtAMR is designed to be flexible, allowing users to choose default settings and accessory files, or using user-supplied criteria which can be tailored as required for local needs. These files are designed to be modifiable by a non-developer (such as a genomics-literate medical microbiologist or scientist), without the requirement for coding skills. This allows users to modify interpretations based on local needs – for example, in a high-TB-burden setting, the threshold for reporting resistance may be lower than a low-TB-burden setting, and as such, the user may choose a lower threshold to report resistance (e.g., report all low-level resistant mutations as resistant). Alternatively, cascade reporting rules may be modified to suit the TB drugs available in different settings. In both situations, we believe it is best for the user to be able to set these parameters, rather than relying on the judgement of the software developer to determine what is ‘best’.

AMR determinants are often reported as a binary ‘present/absent’ variable, inferring a ‘resistant’ or ‘susceptible’ phenotype respectively, but this does not always accurately reflect the complexity of genotype-phenotype correlations. It is important to ensure that mutations used to infer resistance have solid evidence supporting a role as genomic determinants of resistance. The new WHO catalogue and documentation provide some information for mutations which are likely to confer ‘borderline’ resistance such as the rpoB His445Asn variant^15^ and confidence levels which may be used to aid interpretation. The structure of tbtAMR allows us to provide meaningful reports to clinicians (based on high confidence mutations), capturing the complexity of genotype-phenotype relationships, whilst still being able to prospectively monitor and validate additional mutations (mutations from all confidence levels). This is particularly relevant for the newer second-line drugs including bedaquiline, delaminid and linezolid, where a paucity of data (locally and globally) still limits our ability to adequately validate inference of resistance to these drugs for reporting in a CPHL setting. Even for drugs which are well studied, discrepancies can still occur, and further research and iterative validation will only improve the performance of tbtAMR and other tools used by Mtb genomics community, ultimately leading to enhanced patient outcomes.

Performing gDST for Mtb in CPHM settings is a balancing act, weighing direct impacts on patient management on one hand, and pragmatic laboratory management with finite resources in the other. Sequencing processes in a CPHL laboratory are usually dictated by factors such as costs and efficiency. Thus, it is not usually possible for CPHM labs to routinely obtain the sequence depth required to reliably identify high confidence variants below 10% allelic frequency^39^. Furthermore, at lower depth of coverage identification of false positive variants^40^ (Figure 2) could lead to erroneous reporting of gDST. The default settings used in tbtAMR aim to address these considerations, particularly for a CPHM setting, either in high-income country or LMIC. However, these settings are not fixed and may be modified for settings with differing cost and capacity considerations, including research settings.

tbtAMR represents a flexible yet robust and accurate pipeline that leverages the validation undertaken here and other quality studies^22,24,32,33^ to achieve the best predictive power for patient management, and simplified outputs for clinical and public health microbiology. Additionally, the tbtAMR tool has also been implemented with a graphic user interface (GUI) as part of the Centre for Pathogen Genomics (CPG) bioinformatics portal (https://portal.cpg.unimelb.edu.au), which provides access to bioinformatics tools for LMICs and others in resource-limited settings. Whilst this does not alleviate the need for medical microbiologists and scientists in resource-limited settings to develop genomic literacy to be able to optimally use tbtAMR and its results, we hope that providing an accessible GUI with relevant documentation will at least open the door for high TB-burden countries to access our tools.

Our results also highlight areas where further research can enhance our knowledge of AMR determinants in Mtb, with the correlation between AMR determinants and phenotypes with new second-line Mtb drugs being a key priority. Further enhancements of diagnostic processes, such as sequencing directly from samples, will allow us to maximise the benefit of genomic technologies for patient management. Ongoing engagement with stakeholders, including researchers, clinicians and public health officials, ensures that the processes established can be improved and refined where needed to continue to improve patient management and inform timely public health responses.

## Supporting information

supplementary methods and results

example reports

catalogue and criteria for tbtAMR

## Acknowledgements

We thank all the laboratory staff at the Mycobacterial Reference Laboratory, Victorian Infectious Disease Reference Laboratory (VIDRL) and the Microbiological Diagnostic Unit Public Health Laboratory (MDU PHL) who undertook receipt, diagnostics, culture, DNA extraction and sequencing of the material used in this publication; the diagnostic laboratories who diligently provide samples for testing; and the Victorian TB Program, and Department of Health Victoria who fund these services and engage enthusiastically with the laboratories. We also thank the developers and maintainers of the open-source software used and cited in this publication. This work was funded by the Australian National Health and Medical Research Council (NHMRC), Medical Research Futures Fund (FSPGN00049), and Investigator Grant (GNT1196103) to BPH.

## Data sharing

Raw sequence data generated at MDU PHL has been uploaded to the Sequence Read Archive under BioProject PRJNA857537. Accessions for all sequences generated and deposited in public repositories can be found in Supplemental File 1. All results used for interpretation and generation of results, including phenotypic DST results and genomic results used in this study are available in Supplemental File 1 and readily available on GitHub. Further instructions for installing and running tbtamr as well as the complete code base are also available on GitHub (https://github.com/MDU-PHL/tbtamr).

## Author Contributions

Kristy Horan – developed software, study design, data analysis, manuscript writing and editing.

Linda Viberg – performed DNA extractions, reviewed data, edited manuscript

Susan Ballard – contributed to study design, laboratory supervision, edited manuscript

Maria Globan – laboratory supervision, reviewed data, edited manuscript

Wytamma Wirth – developed software, edited manuscript

Katherine Bond – study design, laboratory supervision, edited manuscript

Jessica R Webb – created figures, edited manuscript

Thinley Dorji – created figures, edited manuscript

Deborah Williamson – contributed to study design, edited manuscript

Michelle Sait – performed genomic sequencing, edited manuscript

Ee Laine Tay – reviewed data, edited manuscript

Justin T Denholm – contributed to study design, edited manuscript

Torsten Seemann – study design, software development, validation protocols, edited manuscript

Benjamin P Howden – funding, laboratory supervision, contributed to study design, edited manuscript

Norelle L Sherry – study design, contributed to software development, reporting templates, data analysis, manuscript writing and editing.

## Notes

https://github.com/MDU-PHL/tbtamr

### Competing Interest Statement

The authors have declared no competing interest.

### Summary of Updates

Manuscript was revised to address reviewer comments and suggestions

