## Supplementary material for "Bringing TB genomics to the clinic: A comprehensive pipeline to predict antimicrobial susceptibility from genomic data, validated and accredited to ISO standards": tables

**Table 1 – Antimycobacterial drugs used to treat TB**^2^

| **Type** | **Antimycobacterial drugs** |
| --- | --- |
| **First-line** | Rifampicin, isoniazid, ethambutol, pyrazinamide |
| **Second-line** | |
| **Group A for MDR TB** | Fluoroquinolones (moxifloxacin or levofloxacin), bedaquiline, linezolid |
| **Group B for MDR TB** | Clofazimine, cycloserine or teridazone |
| **Group C for MDR TB** | Delaminid, imipenem-cilastatin or meropenem, ethambutol*, pyrazinamide*, amikacin or streptomycin, ethionamide, para-aminosalicylic acid |
| **Others**  **(Not recommended)** | Kanamycin, capreomycin, gatifloxacin, clavulanic acid |

* where DST indicates susceptibility

**Table 2 – Performance of tbtAMR pipeline to predict phenotypic susceptibility for first- and second-line antimycobacterial drugs**

| **Drug (n)** | **Sensitivity (%,95%CI)** | **Specificity (%,95%CI)** | **PPV (%,95%CI)** | **NPV (%,95%CI)** |
| --- | --- | --- | --- | --- |
| Rifampicin (n = 12,505) | 98.1 (97.6,98.4) | 98.1 (97.7,98.3) | 96.3 (95.7,96.8) | 99.0 (98.8,99.2) |
| Isoniazid (n = 12,545) | 94.2 (93.5,94.9) | 99.1 (98.9,99.3) | 98.5 (98.1,98.8) | 96.5 (96.0,96.9) |
| Ethambutol (n = 10,348) | 90.6 (89.2,91.8) | 95.0 (94.5,95.5) | 81.0 (79.3,82.6) | 97.7 (97.4,98.0) |
| Pyrazinamide (n = 3,174) | 80.8 (76.4,84.5) | 98.8 (98.3,99.1) | 89.2 (85.4,92.2) | 97.6 (96.9,98.1) |
| Bedaquiline (n = 9.339) | 43.2 (33.3,53.6) | 98.2 (97.9,98.5) | 18.5 (13.8,24.4) | 99.5 (99.3,99.6) |
| Linezolid (n = 8,017) | 40.9 (31.2,51.4) | 99.9 (99.8,100.0) | 83.7 (70.0,91.9) | 99.3 (99.1,99.5) |
| Moxifloxacin (n = 7,349) | 93.3 (91.5,94.7) | 97.3 (96.9,97.7) | 84.1 (81.8,86.1) | 98.9 (98.7,99.2) |
| Clofazimine (n = 8,591) | 15.5 (10.3,22.7) | 98.4 (98.1,98.7) | 13.1 (8.6,19.3) | 98.7 (98.4,98.9) |
| Amikacin (n = 10,054) | 88.0 (85.6,90.0) | 98.4 (98.2,98.7) | 83.5 (80.9,85.8) | 98.9 (98.7,99.1) |
| Delamanid (n = 8,882) | 17.0 (10.8,25.9) | 99.9 (99.8,100.0) | 69.6 (49.1,84.4) | 99.1 (98.9,99.3) |
| Ethionamide (n = 9,112) | 86.9 (85.1,88.5) | 94.5 (93.9,95.0) | 74.9 (72.8,76.9) | 97.4 (97.1,97.8) |
| Streptomycin (n = 123) | 95.1 (88.0,98.1) | 78.6 (64.1,88.3) | 89.5 (81.3,94.4) | 89.2 (75.3,95.7) |
| Capreomycin (n = 304) | 92.7 (86.3,96.3) | 98.5 (95.6,99.5) | 97.1 (91.9,99.0) | 96.0 (92.3,98.0) |
| Kanamycin (n= 10,397) | 86.6 (84.4,88.5) | 97.2 (96.8,97.5) | 77.4 (74.9,79.7) | 98.5 (98.2,98.7) |
