## supplementary methods and results for "Bringing TB genomics to the clinic: A comprehensive pipeline to predict antimicrobial susceptibility from genomic data, validated and accredited to ISO standards"

#### Bacterial culture

Clinical samples collected for suspected TB were processed according to routine methods at diagnostic laboratories throughout the state of Victoria, Australia (population 6.71 million in 2022), where routine culture is performed using both broth (mycobacterial growth indicator tubes (MGIT)) and solid culture media. In Victoria, primary samples were cultured both in-house and within external laboratories. Samples with acid-fast bacilli detected were referred to the Mycobacterial Reference Laboratory (MRL) at the Victorian Infectious Diseases Reference Laboratory (VIDRL) for identification and phenotypic susceptibility testing. Broth cultures (MGITs) were sub-cultured onto solid media to provide sufficient material for downstream processes.

*DNA extraction*

DNA was extracted from solid cultures and broth cultures (MGIT) as previously described^1^ with minor modifications. In brief, 3 x 1µl loops of culture were resuspended in 700µL TE and heat killed at 95℃ for 15minutes. For MGITs, 1ml aliquots were heat killed followed by centrifugation and resuspension of the pellet into 700 µL TE. Cells were lysed through mechanical disruption and DNA precipitated with ethanol and sodium acetate followed by elution into EB buffer (QIAGEN).

*Whole genome sequencing (WGS)*

Extracted DNA from solid or broth (MGIT) cultures was transferred to the Microbiological Diagnostic Unit Public Health Laboratory (MDU PHL) for WGS, starting with NexteraXT (Illumina) library preparation according to manufacturer’s instructions, then paired-end short-read sequencing on Illumina NextSeq500/550 platforms.

*Phenotypic DST*

Phenotypic drug susceptibility testing was set up for first line drugs in the BACTEC MGIT 960 system according to WHO guidelines^2^. If resistance at the critical concentrations (rifampicin 0.5µg/mL, isoniazid 0.1µg/mL, ethambutol 5.0µg/mL and pyrazinamide 100µg/mL) was detected, test was repeated and simultaneously second line drugs were set up (amikacin 1.0µg/mL, capreomycin 2.5µg/mL, ethionamide 5.0µg/mL, kanamycin 2.5µg/mL, ofloxacin 2.0µg/mL, moxifloxacin 0.25/1.0µg/mL and isoniazid 0.4µg/mL).

#### tbtAMR design and accessory files

#### Cascade reporting structure

The *db_config.json* file includes a description of the parameters for the mutational catalogue, which drugs to include, reported values/terms for confidence and resistance levels, and a cascade reporting structure. Cascade reporting is a selective reporting strategy where the most appropriate susceptibility results are released based on the resistance of the isolate tested. For example, only results for the first-line TB drugs (plus moxifloxacin, in case of contraindications to other first-line agents) are released for fully susceptible isolates, whilst the full range of results (first, second and third-line agents, where validated) are released for RR-TB and MDR/pre-XDR/XDR-TB isolates. This strategy is used widely in clinical and public health laboratories as an antimicrobial stewardship measure to prevent inappropriate antimicrobial choice, but practices will differ between laboratories (and is not required in research settings), hence rules for reporting can be defined by the user (Supplementary Figure 2).

**Classification rules**

Predicted drug resistance profiles are reported, consistent with the WHO classifications^3^. Pre-extensive drug resistance (pre-XDR) and XDR are reported together, as resistance to bedaquiline and linezolid cannot yet be easily inferred from genomic data.

#### Reverification strategy

It is important to maintain the integrity of any process where the outcome is to be used in informing public health reporting and patient management. Many bioinformatics tools and databases are updated frequently and whilst it may be desirous to always have the most up to date versions, it is also important to ensure that no degradation of quality results occurs as a result. Therefore, it is important to have a robust reverification strategy in place to assess the impact of any changes or updates. Before any updates to the underlying tools of tbtAMR, including mutAMR and its dependencies or new mutations, the potential impact will first need to be assessed.

1. If updates include changes to the methodology used in detection of variants, the performance of tbtAMR to identify accurate sequence will be assessed using the simulated dataset described in this publication.
2. If updates involve changes to the codebase, the performance of tbtAMR to predict AMR will be assessed using sequences from samples submitted to the Victorian MRL.
3. If any modifications to the catalogue are made or to the criteria that determine interpretation and/or classification or reporting of AMR in Mtb (including addition of new drugs for reporting), the performance will be assessed by using sequences from samples submitted to the Victorian MRL.

Any degradation of any performance metric compared to the original validation will be assessed to determine the impact on clinical reporting. Some discordances may result in improvements to prediction of AMR (increases in sensitivity and/or specificity etc), whilst others may result in changes to interpretation, such as level of resistance or confidence in the prediction. In rare cases, updates could lead to a reduction in the confidence in results, such as failure to detect variants and/or incorrect genomic DST result, in which case the updates will be rejected, and the existing versions retained.

Ongoing prospective validation may also be undertaken for new drug combinations using sequences submitted as part of normal Victorian MRL and MDU PHL processes.

### Supplementary Figures and tables

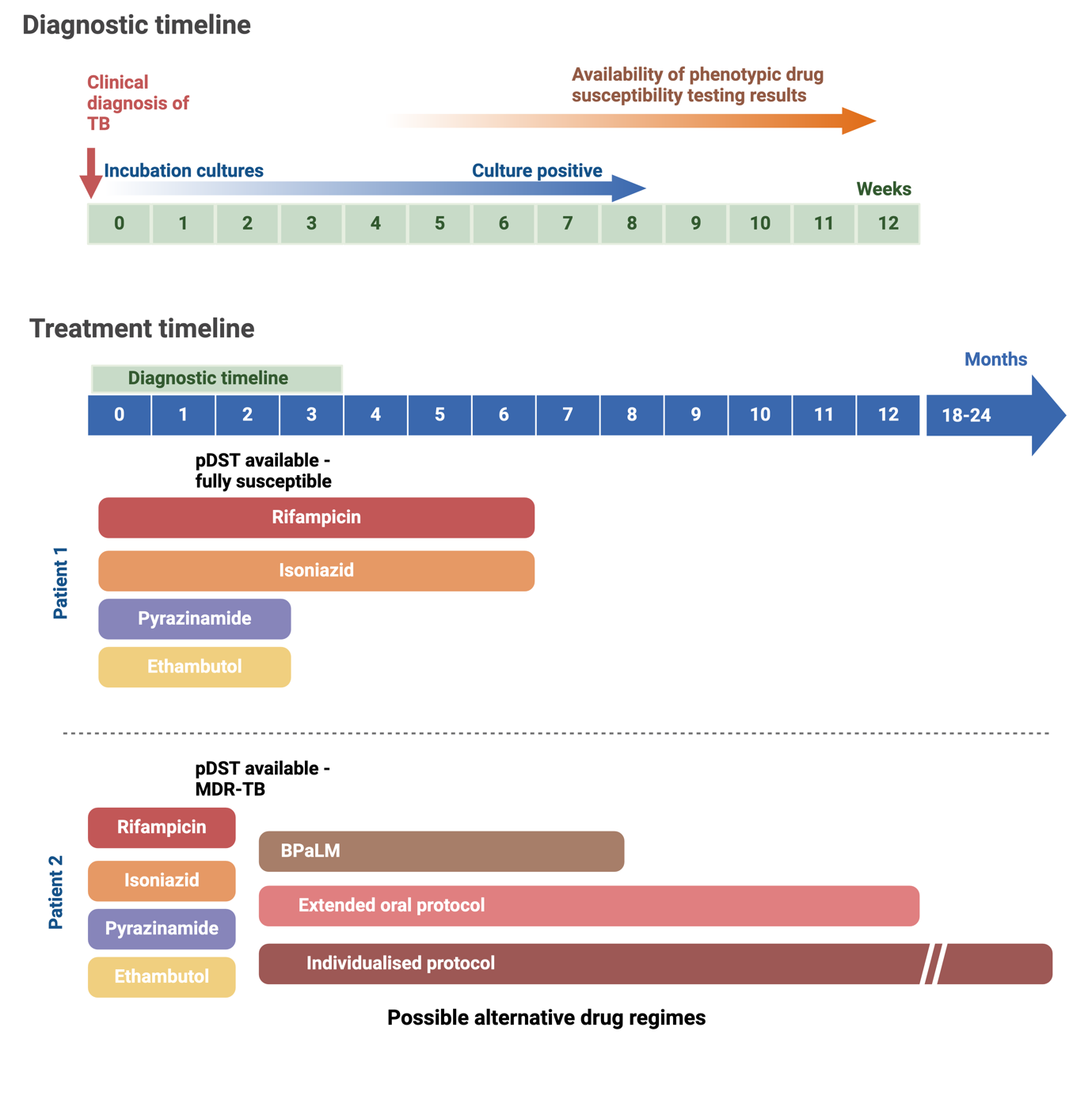

**A**

**B**

#### Supplementary Figure 1 – Outline of common Mtb treatment regimens

**A.** Diagnostic processes for confirmation and determination of drug-resistance for TB can occur over a time scale of weeks, following initial diagnosis, with variations due to laboratory capability and culturability of the samples. Liquid cultures (such as MGITs) are mostly positive in the first 3-4 weeks of culture, whilst longer times to culture positivity are more common with solid cultures and in resource-limited settings. **B.** Upon diagnosis, Mtb is usually treated with four first-line drugs: rifampicin, isoniazid, pyrazinamide and ethambutol. If DST reveals a susceptible phenotype (Patient 1 in Panel B), drug burden may be reduced, such as cessation of ethambutol, and pyrazinamide after the first two months (intensive phase), meaning most patients with drug-susceptible Mtb complete the remaining therapy with only two drugs, rifampicin and isoniazid. Duration depends on the clinical syndrome and is usually 6-12 months. Drug resistance may be identified either when results of pDST are available (Patient 2 Panel B), or earlier if a rapid diagnostic test (e.g. GeneXpert MTB/RIF PCR testing) is available. When likely drug resistance is identified, an alternative treatment regimen is used, with the exact regimen depending on the resistance identified. The BPaLM (bedaquiline, pretomanid, linezolid and moxifloxacin) protocol is a short-duration regimen for MDR or RR-TB taken orally for 6 months. The extended oral protocol includes bedaquiline, moxifloxacin, ethionamide, ethambutol, high dose isoniazid, pyrazinamide and clofazimine for varying overlapping times for 9 months. Other individualised protocols may also be used to address specific resistance profiles and tolerance of the patient. Variations on these timelines are common, depending on timing and type of diagnostic results (e.g. Gene Xpert Mtb/RIF PCR assays providing early results for rifampicin resistance, and whether Mtb culture and DST are routinely performed.

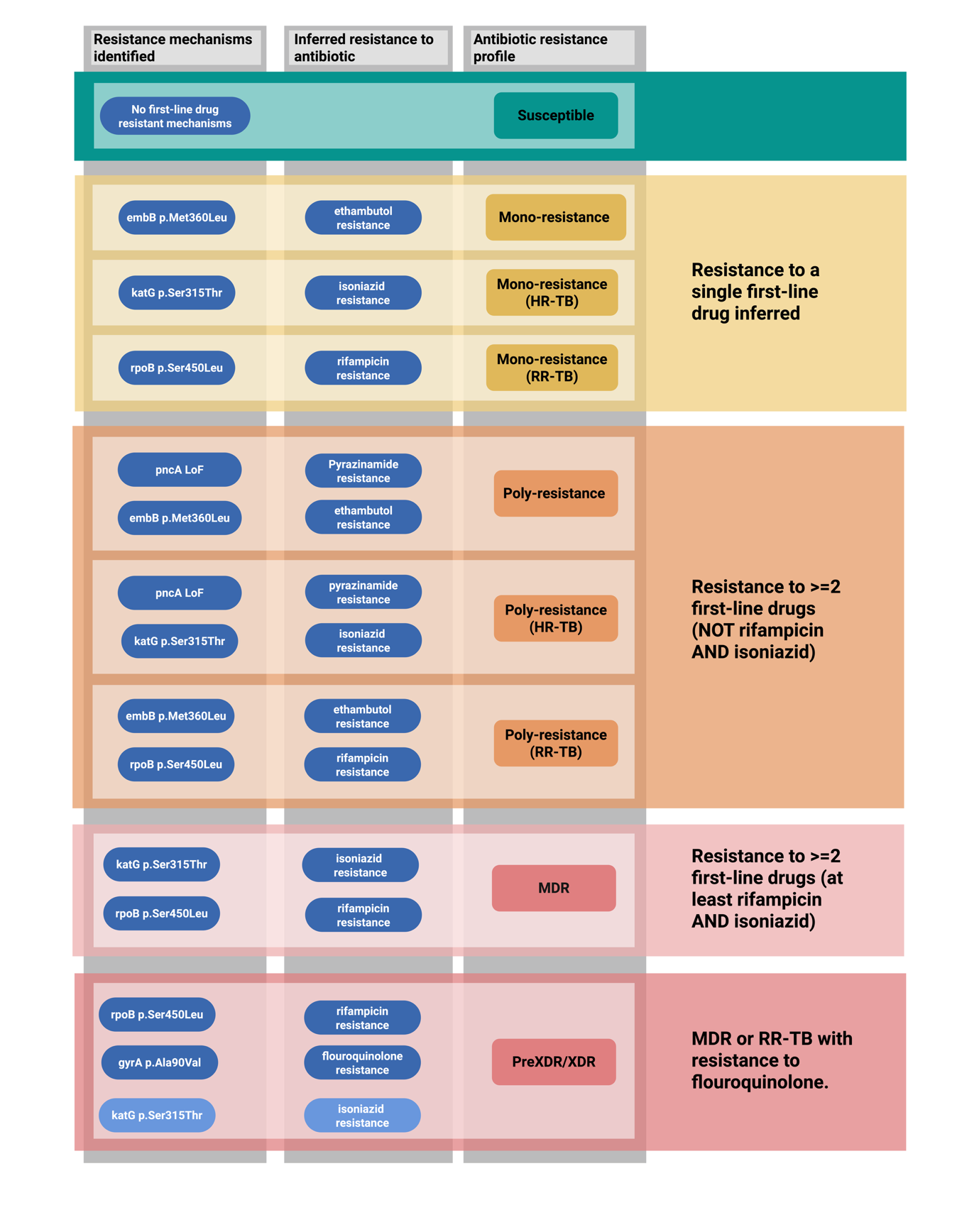

#### Supplementary Figure 2 – Reporting of predicted drug resistance profile

Drug resistant profile is based on the WHO guidelines^3^. Note that Pre-extensive (Pre-XDR) and Extensive Drug Resistance (XDR) cannot yet be differentiated from genomic data (distinguished by bedaquiline and linezolid resistance) and are hence reported as a single group.

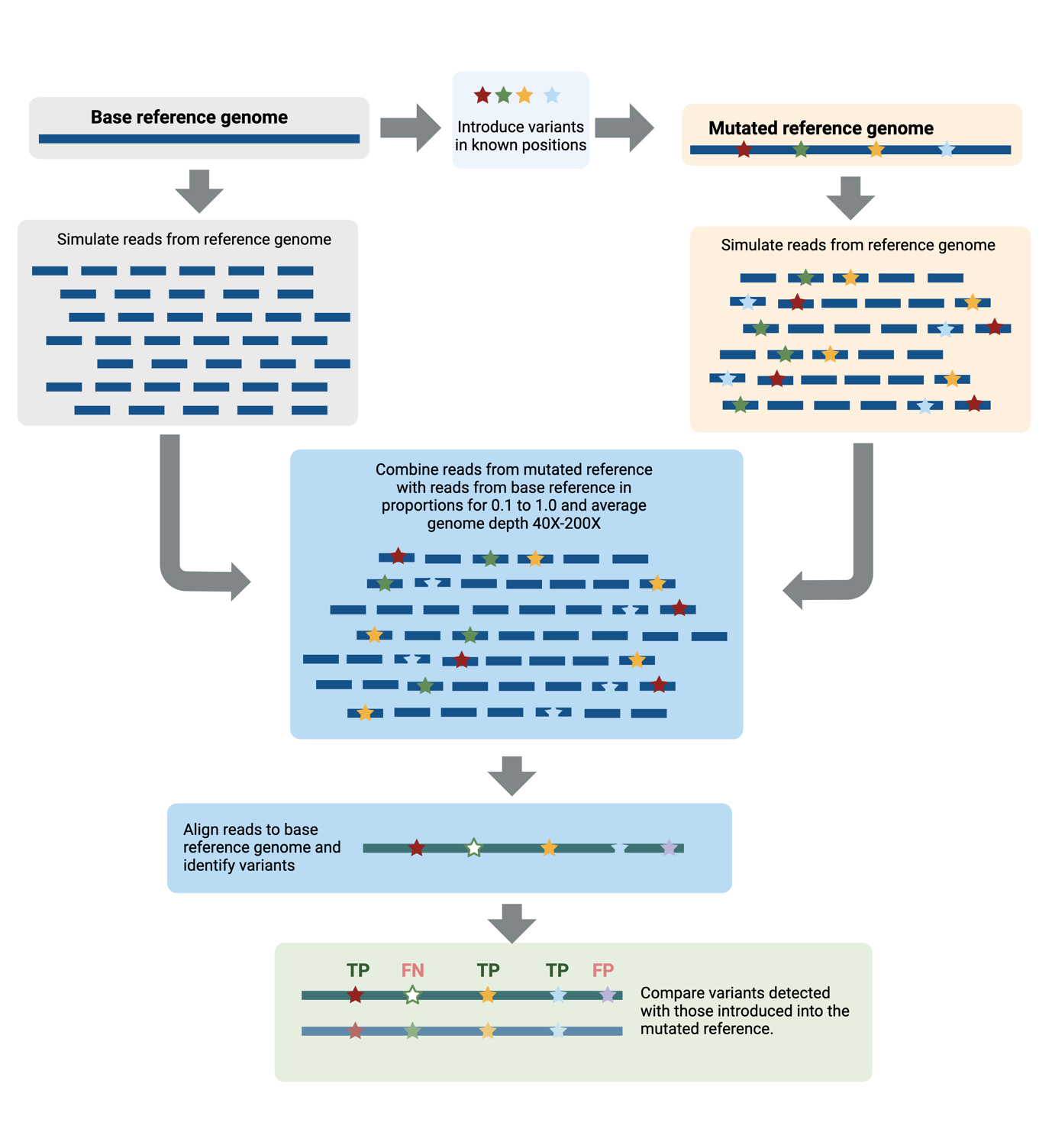

#### Supplementary Figure 3 – Generation and analysis of simulated reads

Simulated reads were generated to mimic the potential mixed alleles observed in Mtb sequencing. Variants were introduced into the complete H37rv reference at known positions and then combined at different proportions across a range of coverages. The simulated sequences (simulated to mimic short-read Illumina sequences) were then analysed using mutamr (implemented in tbtamr to detect variants), and the recovery of the introduced variants was assessed. A true positive (TP) result was observed where an introduced variant was correctly identified; a false positive (FP) result was observed where a variant was reported that was not known to be present; and a false negative (FN) result was observed where an introduced variant was not correctly identified. Mtb, *Mycobacterium tuberculosis.*

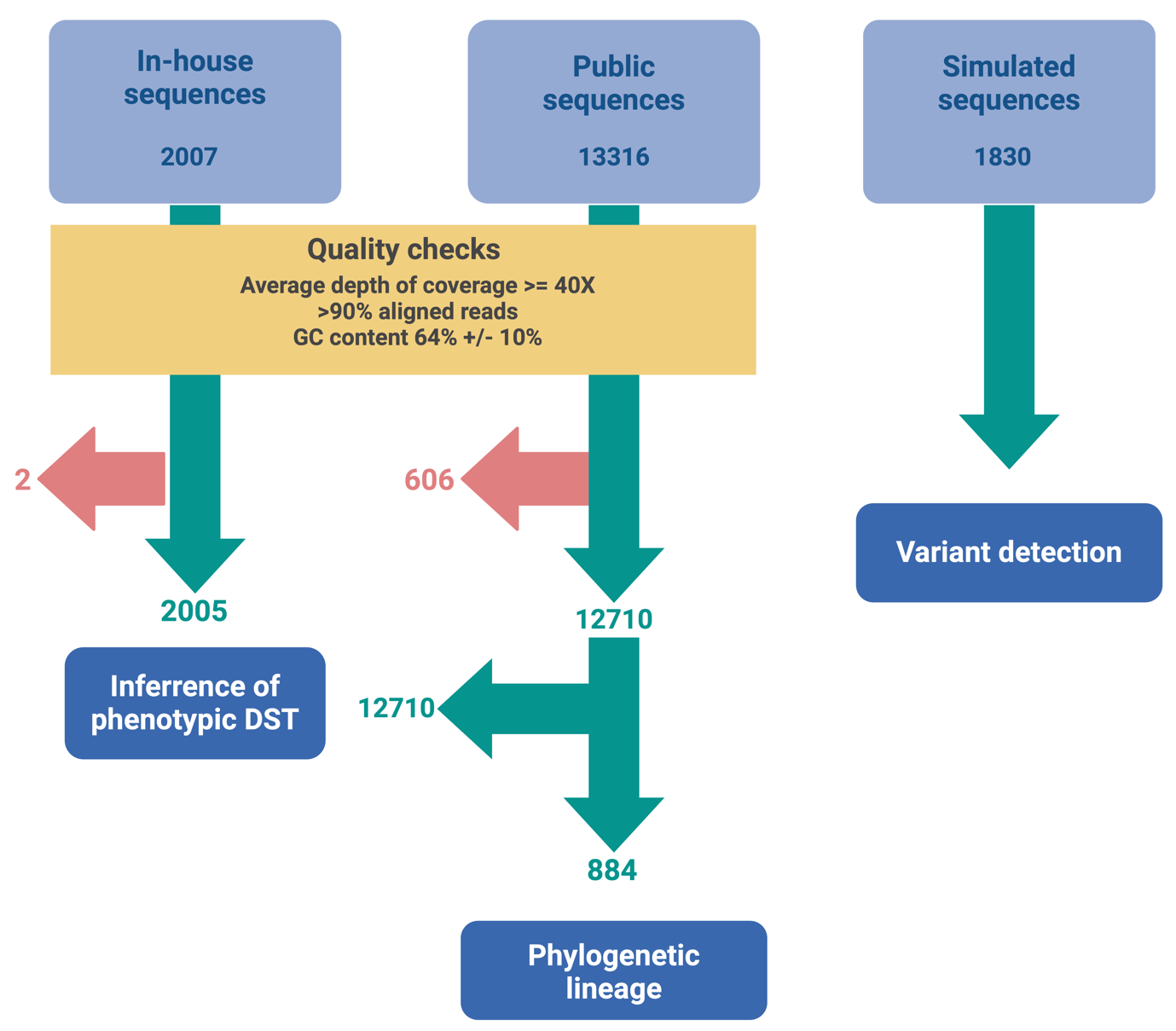

#### Supplementary Figure 4 – Validation dataset

Data generated in-house at MDU, downloaded from public datasets and also simulated data were used to validate variant detection, phylogenetic lineage and genomic DST. Red indicates the number of sequences excluded due to the failure to meet quality requirements.

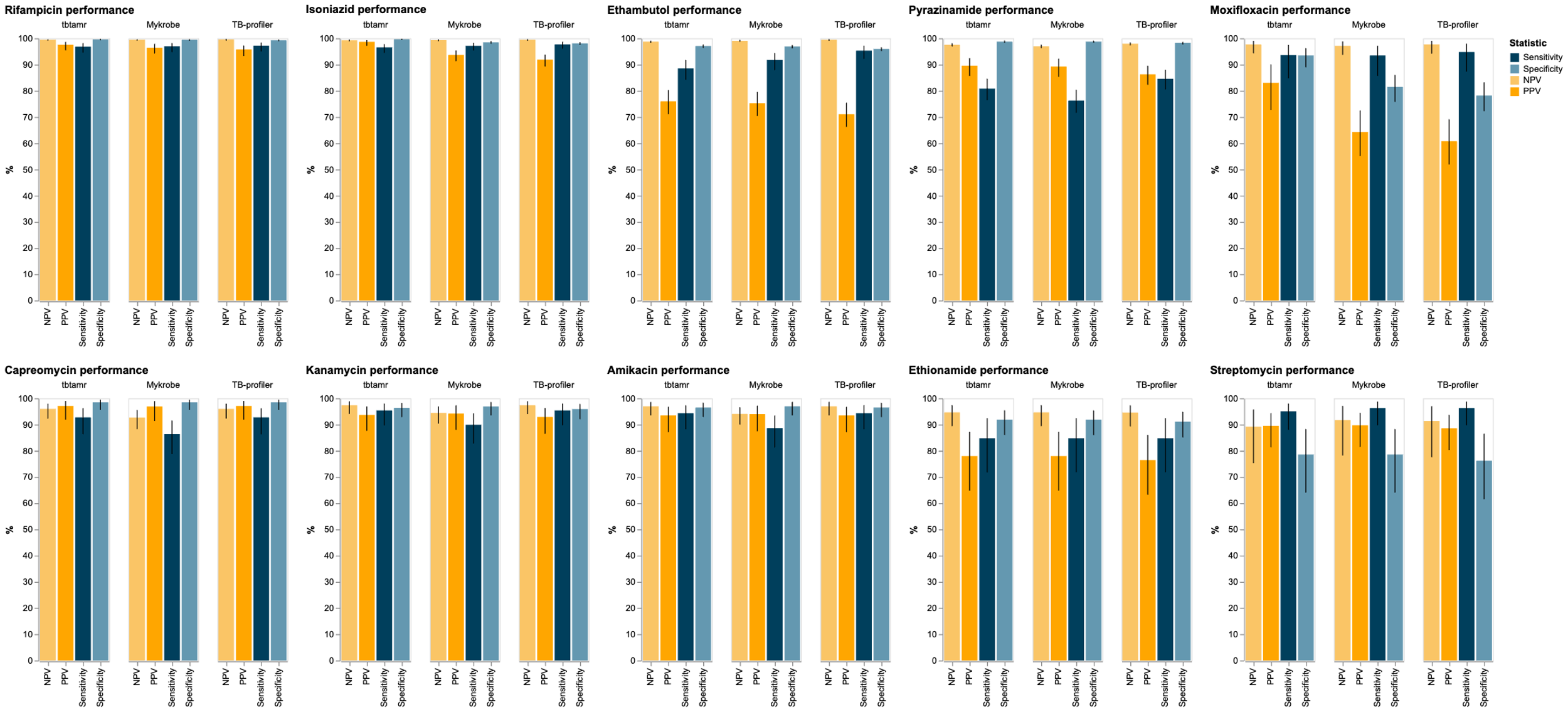

#### Supplementary Figure 5 – Comparsion of tbtAMR, Mykrobe and TB-Profiler performance for inference of phenotypic susceptibility in Mtb

Inferred phenotypes predicted by tbtAMR (v 1.0.3), Mykrobe (v 0.12.1) and tb-profiler (version 6.2.0) using default settings in for all tools were compared to phenotypic DST data, true positive (TP) was recorded when a mutation was reported and the phenotype was resistant; a true negative (TN) was recorded when no mutation was reported and the phenotype was susceptible; a false positive (FP) was recorded when a mutation was reported, but the phenotype was susceptible, and a false negative (FN) was recorded when no mutation was reported, and the phenotype was resistant. Sensitivity, specificity, positive and negative predictive values calculated and expressed as a percentage (error bars indicate 95% confidence intervals). DST, direct susceptibility testing; PPV, positive predictive value; NPV, negative predictive value.

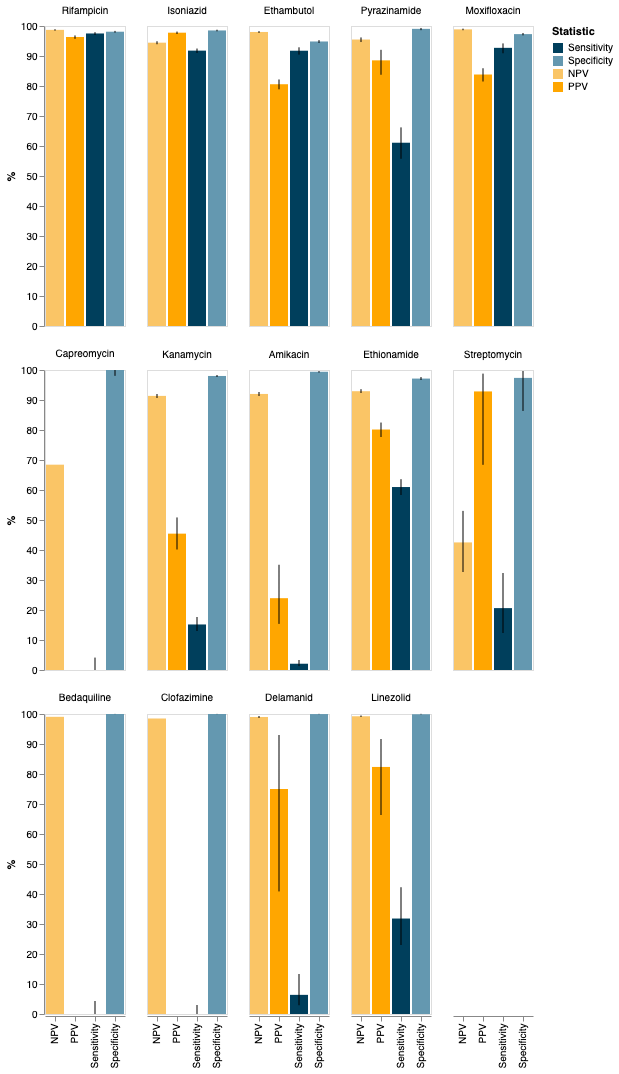

#### Supplementary Figure 6 – Performance of WHO catalogue version 1 run with tbtAMR for inference of phenotypic susceptibility in Mtb

Inferred phenotypes predicted by the tbtAMR pipeline were compared to phenotypic DST data, true positive (TP) was recorded when a mutation was reported and the phenotype was resistant; a true negative (TN) was recorded when no mutation was reported and the phenotype was susceptible; a false positive (FP) was recorded when a mutation was reported, but the phenotype was susceptible, and a false negative (FN) was recorded when no mutation was reported, and the phenotype was resistant. Sensitivity, specificity, positive and negative predictive values calculated and expressed as a percentage (error bars indicate 95% confidence intervals). DST, direct susceptibility testing; PPV, positive predictive value; NPV, negative predictive value.

#### Supplementary Table 1 – Examples of interpretation rules used in tbtAMR

tbtAMR allows users to supply their own rules for interpretation of the impact of a detected variant. Below are descriptions of the types of rules and some examples of how they are used.

| Rule type | Description | Example rules |
| --- | --- | --- |
| Default | - These rules are the base rules for interpreting the impact of detected variants. - A default rule is required for all drugs that the user wants to detect - A user supplies the columns defining the rules and the valid values allowed, as well as the result to apply if these rules are met. | Where drug is rifampicin, if the ‘confidence level’ of variants identified is ‘1) Assoc w R’ or ‘2) Assoc w R – Interim’ then report ‘Resistant’ |
| Override | - These rules address edge cases in interpreting the likely impact of a variant. They are also able to capture more complex interactions between mechanisms. - Override rules will take precedence over default rule. - Similar to default rules, users supply columns and values as well as the result to report if the rule is met. - A simple rule may report a different result to the default rule based on the override rule. - More complex override rules may involve multiple variants and/or multiple columns. | *Simple override rule*   - If there is a single variant that equals rpoB_p.Asp435Tyr, then report ‘low-level resistant’ - this will override the default rule (which is to report Resistant). - if there is a single variant and it is in the *inhA* gene, then report ‘Low-level resistant’ - this will override the default rule (which is to report Resistant).   *Complex override rule*   - If *mmpL5* is not wild-type and Rv0678 has a loss of function (LoF) mutation, stop or frameshift, then report ‘Susceptible’. This will override the default rule (which is to report Resistant). |

#### Supplementary Table 2 – Discordant Phylogenetic lineage

Phylogenetic lineages reported by tbtAMR were compared to those published, with only one of the 13 discordant results not being due the detection of mixed lineages by tbtAMR.

| **Accession** | **Published lineage**^5^ | **tbtAMR lineage** |
| --- | --- | --- |
| ERR067636 | lineage4 | lineage4;lineage2 |
| ERR067732 | lineage4 | lineage2;lineage4 |
| ERR2510523 | lineage3 | lineage4;lineage3 |
| ERR2510554 | lineage1 | lineage1;lineage4 |
| ERR2510576 | lineage3 | lineage3;lineage4 |
| ERR2510682 | lineage2 | lineage2;lineage4 |
| ERR2512421 | lineage3 | lineage4;lineage3 |
| ERR2513452 | lineage4 | lineage2;lineage4 |
| ERR2513557 | lineage1 | lineage1;lineage4 |
| ERR2514865 | lineage1 | La1;lineage1 |
| ERR2515255 | lineage2 | lineage4;lineage2 |
| ERR2515388 | lineage2 | lineage4;lineage2 |
| SRR6339653 | lineage1 | lineage4 |
