## Supplementary material for "Bringing TB genomics to the clinic: A comprehensive pipeline to predict antimicrobial susceptibility from genomic data, validated and accredited to ISO standards": example reports

### Your Laboratory Name

Address
Facsimile: (055) 5555 5555



#### *MYCOBACTERIUM TUBERCULOSIS:*
#### WHOLE GENOME SEQUENCING REPORT

| | | | |
|---|---|---|---|
| To: | Recipient<br>Recipient's Laboratory | Sample ID:<br>Report ID:<br>Issue date: | 2000-123456<br>X123456<br>1-Feb-2000 |

| **Patient details** | | **Submission details** | |
|---|---|---|---|
| Patient name: | FAKE, Name | Submitted item: | 2000-123456 |
| Patient ID | UR3456789 | Primary lab ID: | X123456 |
| Sex / Date of birth: | M / 1-Jan-1986 | Date collected: | 1-Jan-2000 |
| Postcode: | 1000 | | |

#### FINAL RESULTS

**Service**
*M. tuberculosis* whole genome sequencing, analysis and predicted antimicrobial susceptibility

**Result**

| | |
|---|---|
| Identification (WGS) | *Mycobacterium tuberculosis* |
| Phylogenetic lineage | Lineage 4 |

**Predicted antimicrobial susceptibility**

Predicted drug resistance category: No first-line drug-resistance predicted

| Antimicrobial | Resistance mechanism detected | Predicted phenotype | Confidence |
|---|---|---|---|
| Rifampicin | No mechanism identified | Susceptible | |
| Isoniazid | No mechanism identified | Susceptible | |
| Pyrazinamide | No mechanism identified | Susceptible | |
| Ethambutol | No mechanism identified | Susceptible | |
| Moxifloxacin | No mechanism identified | Susceptible | |

Database version A database version

**Comments**
'No mechanism identified' indicated that no resistance-conferring mutation from the reference database was detected in this isolate. Absence of mutations does not exclude phenotypic resistance, and the presence of mutations does not always correlate with phenotypic resistance. Please call the Medical Microbiologist if you wish to discuss further.

**Laboratory accreditation details**

**Report authorised by** A. PATHOLOGIST

#### Your Laboratory Name

Address
Facsimile: (055) 5555 5555


### *MYCOBACTERIUM TUBERCULOSIS:*
### WHOLE GENOME SEQUENCING REPORT

| | | | |
|---|---|---|---|
| To: | Recipient<br>Recipient's Laboratory | Sample ID:<br>Report ID:<br>Issue date: | 2000-123456<br>X123456<br>1-Feb-2000 |

| **Patient details** | | **Submission details** | |
|---|---|---|---|
| Patient name: | FAKE, Name | Submitted item: | 2000-123456 |
| Patient ID | UR3456789 | Primary lab ID: | X123456 |
| Sex / Date of birth: | M / 1-Jan-1986 | Date collected: | 1-Jan-2000 |
| Postcode: | 1000 | | |

#### FINAL RESULTS

**Service**
*M. tuberculosis* whole genome sequencing, analysis and predicted antimicrobial susceptibility

**Result**

| | |
|---|---|
| Identification (WGS) | *Mycobacterium tuberculosis* |
| Phylogenetic lineage | Lineage 2 |

**<u>Predicted antimicrobial susceptibility</u>**

Predicted drug resistance category: Mono-resistance predicted

| Antimicrobial | Resistance mechanism detected | Predicted phenotype | Confidence |
|---|---|---|---|
| Rifampicin | No mechanism identified | Susceptible | |
| Isoniazid | No mechanism identified | Susceptible | |
| Pyrazinamide | pncA p.Ile90Ser | Resistant | High |
| Ethambutol | No mechanism identified | Susceptible | |
| Moxifloxacin | No mechanism identified | Susceptible | |

Database version A database version

**Comments**
”Confidence level” describes the volume of data supporting a mutation conferring a resistant phenotype, based on genotype-phenotype correlations in the WHO Mutation Catalog. ’High confidence’ indicates a strong link (odds ratio (OR) >1) supported by experimental data; ’Moderate confidence’ indicates a strong link (OR >1); and ’Unconfirmed’ indicates a mutation has been observed infrequently, and hence links to phenotypic resistance are unclear. ‘No mechanism identified’ indicated that no resistance-conferring mutation from the reference database was detected in this isolate. Absence of mutations does not exclude phenotypic resistance, and the presence of mutations does not always correlate with phenotypic resistance. Please call the Medical Microbiologist if you wish to discuss further.

**Laboratory accreditation details** **Report authorised by** A. PATHOLOGIST

**Your Laboratory Name**
Address
Facsimile: (055) 5555 5555


### *MYCOBACTERIUM TUBERCULOSIS:* WHOLE GENOME SEQUENCING REPORT

| | | | |
|---|---|---|---|
| To: | Recipient<br>Recipient's Laboratory | Sample ID: | 2000-123456 |
| | | Report ID: | X123456 |
| | | Issue date: | 1-Feb-2000 |

**Patient details**
Patient name: FAKE, Name
Patient ID UR3456789
Sex / Date of birth: M / 1-Jan-1986
Postcode: 1000

**Submission details**
Submitted item: 2000-123456
Primary lab ID: X123456
Date collected: 1-Jan-2000

#### FINAL RESULTS

**Service**
*M. tuberculosis* whole genome sequencing, analysis and predicted antimicrobial susceptibility

**Result**
Identification (WGS) *Mycobacterium tuberculosis*
Phylogenetic lineage Lineage 2

**Predicted antimicrobial susceptibility**

Predicted drug resistance category: Mono-resistance predicted

| Antimicrobial | Resistance mechanism detected | Predicted phenotype | Confidence |
|---|---|---|---|
| Rifampicin | No mechanism identified | Susceptible | |
| Isoniazid | inhA c.-777C>T | Low-level resistant | High |
| Pyrazinamide | No mechanism identified | Susceptible | |
| Ethambutol | No mechanism identified | Susceptible | |
| Moxifloxacin | No mechanism identified | Susceptible | |

Database version A database version

**Comments**
”Confidence level” describes the volume of data supporting a mutation conferring a resistant phenotype, based on genotype-phenotype correlations in the WHO Mutation Catalog. ’High confidence’ indicates a strong link (odds ratio (OR) >1) supported by experimental data; ’Moderate confidence’ indicates a strong link (OR >1); and ’Unconfirmed’ indicates a mutation has been observed infrequently, and hence links to phenotypic resistance are unclear. ‘No mechanism identified’ indicated that no resistance-conferring mutation from the reference database was detected in this isolate. Absence of mutations does not exclude phenotypic resistance, and the presence of mutations does not always correlate with phenotypic resistance. Please call the Medical Microbiologist if you wish to discuss further.

**Laboratory accreditation details**

**Report authorised by** A. PATHOLOGIST

### Your Laboratory Name

Address
Facsimile: (055) 5555 5555



#### *MYCOBACTERIUM TUBERCULOSIS:*
#### WHOLE GENOME SEQUENCING REPORT

| | | | |
|---|---|---|---|
| To: | Recipient<br>Recipient's Laboratory | Sample ID:<br>Report ID:<br>Issue date: | 2000-123456<br>X123456<br>1-Feb-2000 |

| **Patient details** | | **Submission details** | |
|---|---|---|---|
| Patient name: | FAKE, Name | Submitted item: | 2000-123456 |
| Patient ID | UR3456789 | Primary lab ID: | X123456 |
| Sex / Date of birth: | M / 1-Jan-1986 | Date collected: | 1-Jan-2000 |
| Postcode: | 1000 | | |

#### FINAL RESULTS

**Service**
*M. tuberculosis* whole genome sequencing, analysis and predicted antimicrobial susceptibility

**Result**
Identification (WGS) *Mycobacterium tuberculosis*
Phylogenetic lineage Lineage 1

**Predicted antimicrobial susceptibility**

Predicted drug resistance category: Multi-drug resistance predicted (MDR-TB)

| Antimicrobial | Resistance mechanism detected | Predicted phenotype | Confidence |
|---|---|---|---|
| Rifampicin | rpoB p.Ser450Leu | Resistant | High |
| Isoniazid | katG p.Ser315Thr | Resistant | High |
| Pyrazinamide | pncA p.Ala102Thr | Resistant | Moderate |
| Ethambutol | No mechanism identified | Susceptible | |
| Moxifloxacin | No mechanism identified | Susceptible | |
| Amikacin | No mechanism identified | Susceptible | |
| Ethionamide | No mechanism identified | Susceptible | |

Database version A database version

**Comments**
'No mechanism identified' indicated that no resistance-conferring mutation from the reference database was detected in this isolate. Absence of mutations does not exclude phenotypic resistance, and the presence of mutations does not always correlate with phenotypic resistance. Please call the Medical Microbiologist if you wish to discuss further.
"Confidence level" describes the volume of data supporting a mutation conferring a resistant phenotype, based on genotype-phenotype correlations in the WHO Mutation Catalog. 'High confidence' indicates a strong link (odds ratio (OR) >1) supported by experimental data; 'Moderate confidence' indicates a strong link (OR >1); and Correlation with phenotypic testing is recommended. Please call the Medical Microbiologist if you wish to discuss further.

**Laboratory accreditation details** **Report authorised by** A. PATHOLOGIST

### Your Laboratory Name

Address
Facsimile: (055) 5555 5555



#### *MYCOBACTERIUM TUBERCULOSIS:* WHOLE GENOME SEQUENCING REPORT

| | | | |
|---|---|---|---|
| To: | Recipient<br>Recipient's Laboratory | Sample ID:<br>Report ID:<br>Issue date: | 2000-123456<br>X123456<br>1-Feb-2000 |

| **Patient details** | | **Submission details** | |
|---|---|---|---|
| Patient name: | FAKE, Name | Submitted item: | 2000-123456 |
| Patient ID | UR3456789 | Primary lab ID: | X123456 |
| Sex / Date of birth: | M / 1-Jan-1986 | Date collected: | 1-Jan-2000 |
| Postcode: | 1000 | | |

#### FINAL RESULTS

**Service**
*M. tuberculosis* whole genome sequencing, analysis and predicted antimicrobial susceptibility

**Result**

| | |
|---|---|
| Identification (WGS) | *Mycobacterium tuberculosis* |
| Phylogenetic lineage | Lineage 3 |

**Predicted antimicrobial susceptibility**

Predicted drug resistance category: Pre-Extensive/Extensive drug-resistance predicted

| Antimicrobial | Resistance mechanism detected | Predicted phenotype | Confidence |
|---|---|---|---|
| Rifampicin | rpoB p.Ser450Leu | Resistant | High |
| Isoniazid | inhA c.-154G>A;katG p.Ser315Thr | Resistant | High |
| Pyrazinamide | pncA p.Ala102Thr | Resistant | Moderate |
| Ethambutol | No mechanism identified | Susceptible | |
| Moxifloxacin | gyrA p.Asp94Gly | Resistant | High |
| Amikacin | No mechanism identified | Susceptible | |
| Ethionamide | inhA c.-154G>A | Resistant | High |

Database version A database version

**Comments**
Genotyping cannot currently distinguish between multidrug resistant TB (MDR TB) and extensively drug-resistant TB (XDR TB), due to the lack of genotypic data for bedaquiline and linezolid resistance.
'No mechanism identified' indicated that no resistance-conferring mutation from the reference database was detected in this isolate. Absence of mutations does not exclude phenotypic resistance, and the presence of mutations does not always correlate with phenotypic resistance. Please call the Medical Microbiologist if you wish to discuss further.
"Confidence level" describes the volume of data supporting a mutation conferring a resistant phenotype, based on genotype-phenotype correlations in the WHO Mutation Catalog. 'High confidence' indicates a strong link (odds ratio (OR) >1) supported by experimental data; 'Moderate confidence' indicates a strong link (OR >1); and Correlation with phenotypic testing is recommended. Please call the Medical Microbiologist if you wish to discuss further.
inhA promoter mutations were previously reported as fabG1

**Laboratory accreditation details**

**Report authorised by** A. PATHOLOGIST
